# AutoSTED: An automated workflow for STED super-resolution imaging of cell nuclei

**DOI:** 10.64898/2025.12.11.693650

**Authors:** Dorian Leger, Milena Ivanisevic, Péter Lénárt

## Abstract

The traditional microscopy workflow involves manual selection of objects one after the other. This is not only time consuming, but inevitably introduces human bias. Here, we describe a workflow, ‘AutoSTED’ for identification of cell nuclei on wide-field fluorescence images and subsequent automated imaging of these nuclei at super-resolution by Stimulated Emission Depletion (STED) microscopy. We demonstrate that in an overnight run we are able to image hundreds of cells selected without bias. The resolution afforded by STED allowed straightforward identification and counting of individual nuclear pore complexes (NPCs), and enabled us to automatically extract parameters such as NPC density under different experimental conditions. Together, we illustrate how automation enables optimal use of instrument time to produce large unbiased datasets.

## Introduction

Traditionally, microscopic samples are analyzed by expert operators who visually screen slides and manually select cells or other structures of interest for imaging and documentation. Although automated whole-slide scanning has become common in histopathology and fluorescence imaging for routine workflows, manual region selection remains the norm for advanced techniques such as super-resolution microscopy (Rimon and Schuldiner, 2011; Boyang et al., 2023).

A major obstacle to full automation is acquisition speed. Super-resolution methods such as Stimulated Emission Depletion (STED) microscopy and Single Molecule Localization Microscopy (SMLM), including PALM and STORM, dramatically improve resolution beyond the diffraction limit but do so at the cost of high illumination intensity, low photon counts per pixel, and serial scanning (Bond et al., 2022). In STED microscopy, pixels as small as 15×15 nm are recorded sequentially, resulting in acquisition times several orders of magnitude longer than those of wide-field fluorescence imaging. Consequently, operators typically spend several minutes acquiring a single image, making manual cell selection an inefficient use of both personnel and instrument time.

Given that a single slide may contain thousands of fluorescently labeled cells, only a small subset— often fewer than a dozen—is typically imaged in detail. This restricted sample size and the inherent subjectivity of manual selection introduce biases and limit statistical robustness. Automated, objective, and high-throughput acquisition workflows are therefore essential to improve reproducibility in super-resolution experiments (Chen et al., 2023).

Here, we present a workflow for automated acquisition on a hybrid wide-field and STED imaging system. Although tailored to our instrument configuration, the approach addresses general challenges in automated super-resolution microscopy. First, wide-field fluorescence images are acquired over an entire slide region, either as a grid or from pre-selected fields of view. Automated algorithms then detect objects of interest, such as cell nuclei, which subsequently trigger higher-resolution STED acquisition. The system determines optimal z-position, automatically focuses, and executes imaging cycles automatically until all identified regions are processed. This approach maximizes information yield while minimizing both acquisition time and photo-damage.

To demonstrate this pipeline, we implemented automated identification of cell nuclei expressing endogenously tagged nuclear pore complexes (NPCs). We imaged NPCs in Nup96-SNAP labeled U2OS cells to quantify NPC density within the nuclear envelope in live cells. These proof-of-principle experiments illustrate the power of automated STED microscopy: individual NPCs, unresolved in confocal or wide-field modes, were clearly resolved and quantified in hundreds of automatically acquired images. The resulting dataset is both large and unbiased, enabling statistically meaningful analysis, supporting reproducible imaging-based research.

## Results

### AutoSTED workflow for automated object identification and STED imaging

Our microscope system is composed of a motorized Nikon Ti2 inverted microscope body with a Hamamatsu ORCA Fusion sCMOS camera mounted on the left camera port for wide-field fluorescence imaging. For STED imaging, a Stedycon scanhead is mounted on the right camera port. The microscope and the sCMOS camera are operated through the NIS-Elements software (Nikon Instruments), the Stedycon is controlled via its own software run in a web browser (Chrome), with all software installed on the same desktop computer (Fig. S1).

While we used proprietary software for hardware control, we designed our integrated workflow in a modular manner and used standard formats and open-source software (Fig. 1A). This should allow the modules to be easily adapted to diverse hardware configurations. The first module transfers, in a standard tiff-format, the wide-field fluorescence image acquired by NIS-Elements software to ImageJ/FIJI (Schindelin et al., 2012). Objects of interest are then identified in this environment and XY coordinates of center of mass of identified objects are written in a CSV file. Optionally, a specific Z coordinate of the object can also be detected, for instance, the position corresponding to the bottom of nucleus of cells. Subsequently, coordinates in the csv file are read by NIS-Elements and the stage is moved to the first XY(Z) position. Then an analog trigger (TTL) is sent to the Stedycon through a NIDAQ card to initiate the acquisition of the STED image.

**Figure 1.**
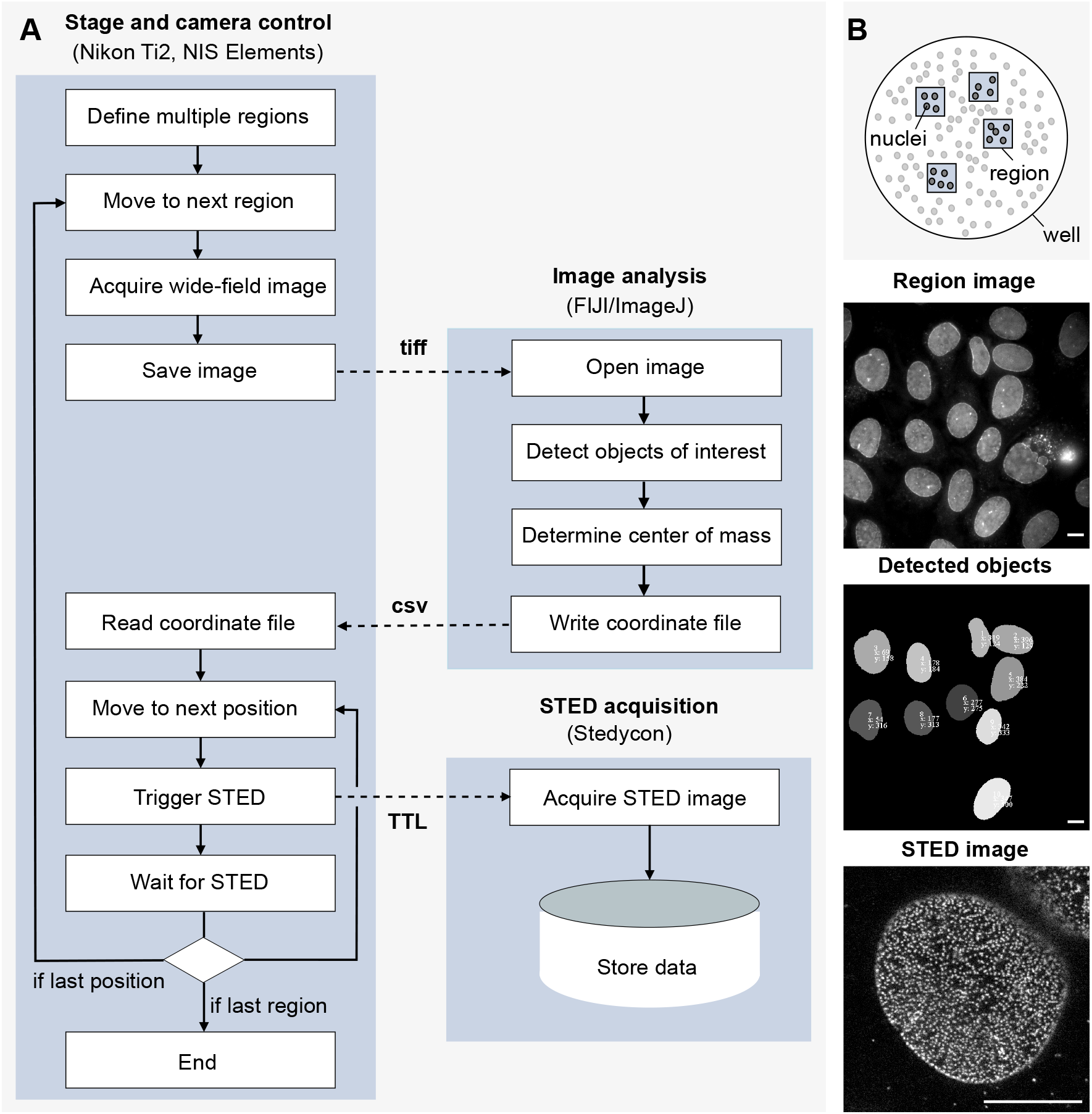
Outline of the AutoSTED adaptive feedback microscopy workflow. **(A)** Flow diagram of the AutoSTED workflow. **(B)** Schematics of the live-cell samples. U2OS cells were grown on 10-well plates. Examples images are shown, (*top*) wide-field fluorescence image of the region with several nuclei stained, (*middle*) label masks of the detected nuclei, (*bottom*) STED image of a single nucleus with the NPCs labeled with SNAP-647-SiR. Scale bars are 10 µm.

To initiate an experiment, the user first marks the regions to be imaged using one of the fluorescent channels and the camera in live mode controlled by the NIS-Elements software (Fig. 1B). Alternatively, a grid of regions can be generated automatically. Then the fluorescence channels are selected again in NIS-Elements to image the overview regions. Next, the user selects the appropriate adaptive microscopy module: for example, the module for detection of labelled nuclei in the XY plane. Then, the user selects STED settings, such as size of ROI, channels, line accumulation, and excitation and depletion laser intensities.

Following these manual steps of initial setup, the user runs the NIS-Elements macros and ImageJ/FIJI macros for automated acquisition. NIS-Elements moves the stage to the first region, captures a wide-field image, saves the image and notifies ImageJ/FIJI that the acquisition is finished. ImageJ/FIJI waits for the notification and then opens the wide-field image of the first region (Fig. 1B). ImageJ/FIJI performs an analysis and writes the coordinates of objects of interest into a CSV file. ImageJ/FIJI then notifies NIS-Elements that the adaptive analysis phase is finished. Next, NIS-Elements reads the CSV file, moves the stage to the first coordinate in the file, and sends an analogue signal to Stedycon to trigger a STED acquisition. NIS-Elements then waits for a designated time and subsequently moves to the coordinates of the next position in the CSV file. When all positions in the CSV file are acquired, NIS-Elements moves the stage to the next region and repeats the loop. When all regions are completed, NIS-Elements ends the experiment, unless a time-loop is set.

Images are stored during the experiment to the local hard drive of the computer. The raw wide-field image of each region and the labelled wide-field image (Fig. 1B), which show the detected objects, are also stored to the local drive of the computer. Prior to the acquisition of a STED image, NIS-Elements makes an entry in a log file which records XYZT position of the image. For multi-well experiments, the log file also indicates the well number.

### Use case: Automated measurement of nuclear pore density live cells with AutoSTED

Nuclear pore complexes (NPCs) are the channels mediating traffic between the cytoplasm and nucleus (Beck and Hurt, 2017; Petrovic et al., 2025). Therefore, the number and density of NPCs are a key reporter of the physiological state of the cell: NPC number is correlated with cell growth, chromatin organization and changes in gene expression. Abnormal NPC number or density is indicative of several pathological conditions, including mis-regulated growth in cancer (Coyne and Rothstein, 2022).

NPCs are one of the largest protein complexes in the cell with approximately 120 nm diameter, so large that individual complexes are visible by fluorescence light microscopy. However, when NPCs are close to one another, conventional light microscopy is not sufficient to resolve, and thus to reliably count individual NPCs (Thevathasan et al., 2019). Reaching a resolution down to 20-30 nm, STED can be used to resolve even NPC substructures; however, this requires relatively high powers of the STED depletion laser. Here we used low powers of STED depletion laser, resulting in lower resolution of approximately 50-60 nm. This resolution is still sufficient to clearly resolve individual NPCs and significantly reduces photo-damage, making it well-suited for live-cell imaging applications. To ensure that we acquire the ‘best-focused’ section, we recorded 5 z-sections 300 nm apart (Fig. 2A). For visualization, we aligned these slices and rendered maximum intensity projections (Fig. 2B).

**Figure 2.**
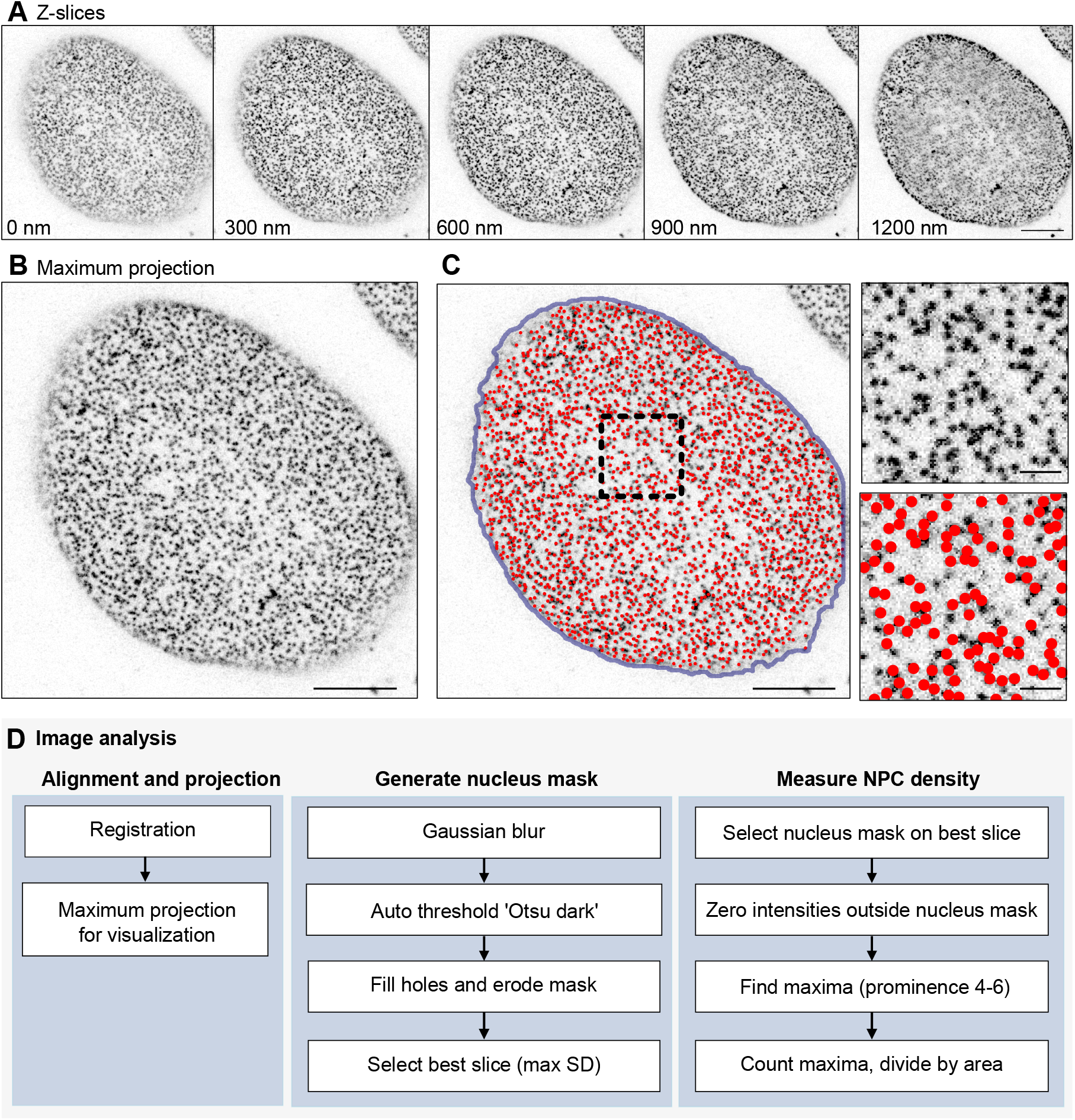
Example of the STED super-resolution recording and steps of image analysis. **(A)** The complete dataset composed of 5 Z-sections of a nucleus labeled with Nup96-SNAP-647-SiR. **(B)** Maximum intensity projection of (A) after alignment of the Z-sections. **(C)** Segmentation of the nucleus (blue outline) and identification of individual NPCs (red circles). Region marked with dashed rectangle is shown at a larger size on the right. **(D)** Flow diagram of the image analysis workflow. Scale bars are 5 µm, and 1 µm for the insets.

The increased resolution of STED enabled a straightforward analysis of the NPC density, by automated counting of NPCS per unit nucleus surface area, in a large number of cells. For this purpose, we assembled a script using standard algorithms available in ImageJ/FIJI (Fig. 2D). After blurring fine structures (Gaussian blur with sigma 7), the area of the nucleus was identified by a simple thresholding step (using Otsu’s method). The resulting mask was then processed further by filling holes and erosion. The standard deviation of intensities within this mask was then used to select the ‘best-focused’ section. Next, NPCs were identified and counted in the nuclear region by using the ‘Find Maxima’ function (using a prominence value between 4 and 6). Figure 2C shows the identified pores overlaid on the maximum projection of the image stack. We manually evaluated the accuracy of this workflow and found that it reliably identified NPCs on the vast majority of acquired images. Additionally, we established criteria, based on which we excluded poor quality images. These included images in which the initial thresholding of the nuclear area failed and the NPC density value fell below 1 or exceeded 10 per µm^2^.

To validate the workflow in a realistic experimental setting, we followed up on previous observations made in HeLa cells (Maeshima et al., 2010). As a cell proceeds through the cell cycle, the size of the nucleus (the surface area) as well and the number of NPCs approximately doubles, maintaining NPC density roughly constant. The study by Maeshima and colleagues shows that formation of new NPCs requires the activity of cyclin-dependent kinases (Cdks), while nuclear growth is independent of Cdk activity. Consequently, inhibition of Cdks leads to a reduction of NPC density in the G2 phase of the cell cycle (Maeshima et al., 2010).

Using our automated workflow, we wanted to confirm findings made earlier in HeLa cells in our U2OS cell line. U2OS cells were synchronized in S-phase by thymidine treatment (Gobran et al., 2025) and were then released and treated with different Cdk inhibitors. First we tested the Cdk1/2 inhibitor Roscovitine (Meijer et al., 1997), which has been used in the aforementioned study (Maeshima et al., 2010). In addition, we used NU-6140, an inhibitor specific to Cdk2 (Talapati et al., 2021), and RO-3306, a commonly used small-molecule inhibitor that is specific to Cdk1 (Vassilev et al., 2006). In these experiments, we could confirm that Roscovitine treatment led to a significant reduction of pore density (Fig. 3A, B). Further, we observed a slightly weaker, but significant reduction in pore density using the Cdk2 inhibitor NU-6140, while cells treated with RO-3306 showed no significant change in NPC density (Fig. 3A, B).

**Figure 3.**
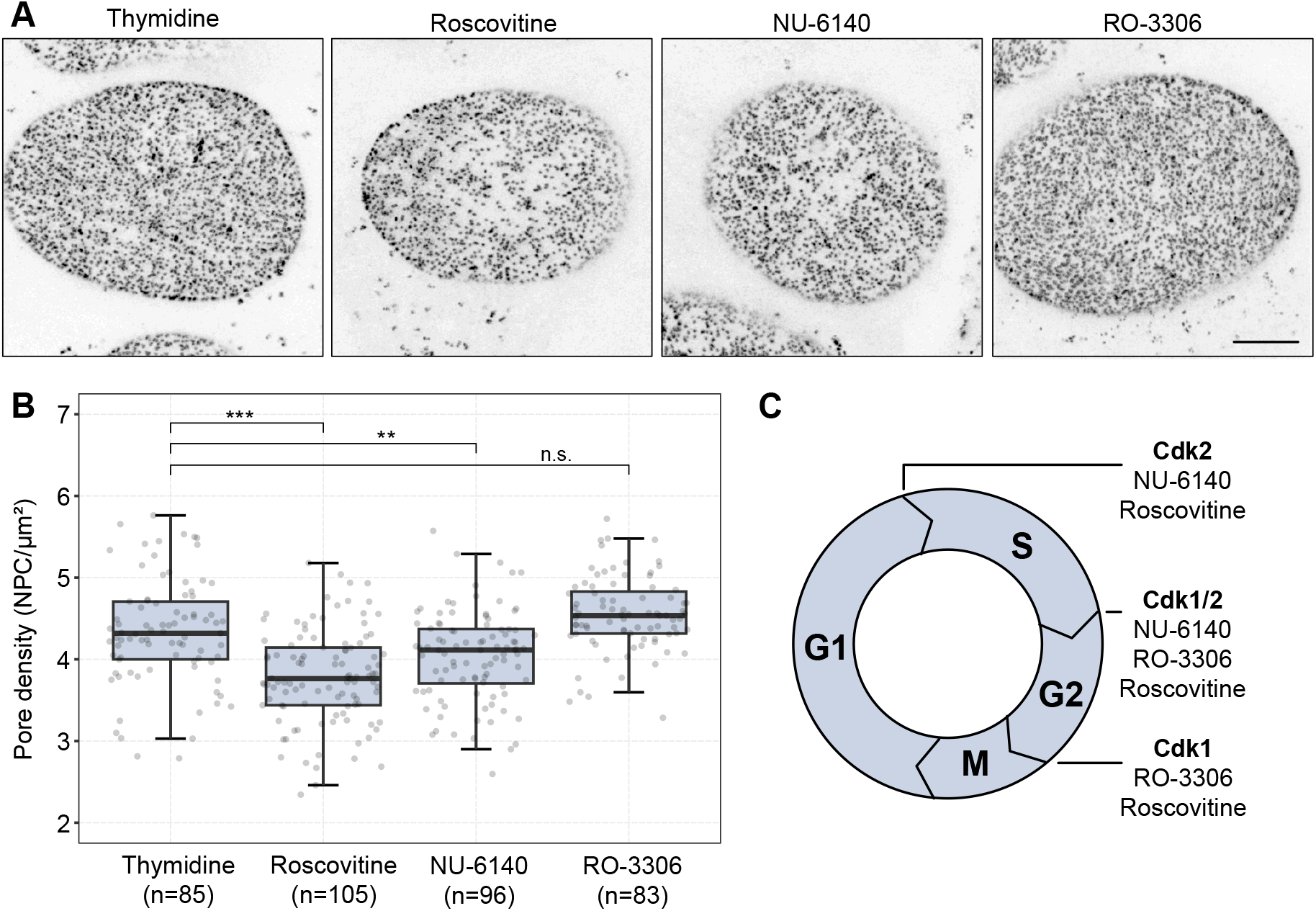
Use case of the AutoSTED workflow. **(A)** Examples of STED images (maximum intensity projections) of cell nuclei labeled with Nup96-SNAP-647-SiR under the indicated experimental conditions. **(B)** Analysis of a complete dataset from an overnight recording. Gray dots show the NPC density of an individual cell, box plots show mean and distribution. n is the number of cells analyzed for the respective treatment. Statistical comparisons were done using Student’s t-test. **(C)** Schematics of cell cycle progression, its control by Cdks, and the respective inhibitors used in the experiments.

These data validate our workflow and demonstrate that it can be used to characterize cells under multiple experimental conditions in a single overnight run, and generate a large and unbiased dataset. While these interpretations need to be further validated, our data confirm that Cdk activity is required for formation of new NPCs, and suggest that this is specifically mediated by the activity of the S-phase Cdk, Cdk2 (typically complexed with Cyclin A), rather than the M-phase kinase, Cdk1-Cyclin B.

## Discussion

In summary, our automated workflow uses gentle wide-field imaging of large fields of cells to identify cell nuclei for STED imaging. We show that this workflow is capable of imaging hundreds of nuclei in an overnight run with a resolution beyond the diffraction limit, allowing counting of NPCs and thus quantification of NPC density. Thereby, the workflow is able to deliver a large and unbiased dataset of multiple experimental conditions, overcoming the inherent limitation of long acquisition times that are typically associated with super-resolution imaging techniques.

While our setup is based on a specific combination of hardware and software components, it serves as a proof-of-concept for the approach, demonstrating its feasibility and potential for future advancements. This is greatly facilitated by the modular design of our workflow, which leverages open-source packages and standard data formats (tiff images, CSV files for coordinates, and TTL triggers). For example, we use NIS-Elements for controlling the microscope and for wide-field image acquisition, but it could be easily replaced by other commercial software or a microscope system controlled by the open source package, MicroManager (Edelstein et al., 2010). Similarly, for high-/super-resolution acquisition, any modality and instrument that can be triggered by a standard TTL signal can be used.

In this example, we used a very simple method to detect objects, i.e. to identify nuclei. While it was sufficient for our application, more complex analysis modules could instead be implemented. In particular, machine-learning and deep-learning-based segmentation tools are revolutionizing image analysis (Celik and Inik, 2026) and could be trained to recognize specific morphologies and thus very specific cellular stages – even without requiring a stage-specific staining. Although these algorithms require substantial computational capacity, it is now feasible to run them on a standard workstation equipped with appropriate GPU to analyze images on-the-fly in an adaptive feedback microscopy workflow. Several of the software tools can be directly integrated with ImageJ/FIJI (e.g. Ilastik, Berg et al., 2019) or the workflow could alternatively be implemented in a Python environment.

In conclusion, our workflow integrates automated object detection with automated microscope control to move the stage to and acquire detected objects without supervision. This enables unsupervised high-throughput experiments with specificity in data acquisition. We demonstrate that such adaptive feedback microscopy workflow is feasible specifically in combination with super-resolution techniques, providing a solution to a major bottleneck caused by slow data acquisition associated with these methods.

## Materials and methods

### Cell culture, reagents and drug treatments

We used the osteosarcoma-derived U2OS cells containing an endogenously tagged SNAP-Nup96 gene (U2OS SNAP-Nup96) (Thevathasan et al., 2019). Cells were grown in high glucose DMEM (Thermo Fisher) supplemented with 10% inactivated fetal bovine serum (FBS), 1% of 100 mM sodium pyruvate (Thermo Fisher) and 1% of 200 mM L-Glutamine (Thermo Fisher). Cells were grown at 37°C in 5% CO_2_, atmospheric O_2_ and saturating humidity.

Cells were plated into 10-well glass bottom imaging plates (Greiner, 543079) or 8-well glass bottom imaging plates (Ibidi, 80807). Two hours before imaging, the culture medium was replaced by imaging medium: FluoroBrite DMEM (Thermo Fisher) containing 10% FBS, 1% of 100 mM sodium pyruvate and 1% of 200 mM L-glutamine, a final concentration of 1 µM verapamil (Spirochrome) and 50 nM of SNAP-647 SiR stain (Lukinavičius et al., 2015). Before imaging, cells were washed twice with 300 µl of pre-warmed imaging medium to remove the excess SNAP dye.

Cells were synchronized in S-phase by thymidine treatment (Gobran et al., 2025). 24 hours after seeding, cells were blocked for 24 hours by adding 2 mM thymidine (Sigma) to the growth medium. Cells were released by exchange of growth medium with imaging medium. After thymidine release, cells were treated with Cdk inhibitors at the following final concentrations: 50 µM Roscovitine (Sigma), 10 µM NU-6140 (Sigma) and 10 µM RO-3306 (Sigma) combined with thymidine. For controls, cells were treated with 2 mM thymidine and equal amount of DMSO. After 4 hours, cells were stained with 50 nM SNAP-Cell 647-SiR (New England Biolabs) and 1 µM Verapamil. After 2 hours of incubation at 37°C, cells were imaged for 15 hours.

### Imaging

Imaging was done on a Nikon Eclipse Ti2 inverted microscope (Nikon Instruments) system with a Hamamatsu ORCA-Fusion camera (Hamamatsu) mounted on the left camera port, and a Stedycon STED scan head (Abberior Instruments) on the right side. For wide-field illumination a Lumencor Sola SE II LED light source was used, and the Stedycon module is equipped with 405, 488, 561 and 640 nm pulsed lasers (except 405 nm) and a 775 nm pulsed STED depletion laser. The microscope is equipped with a motorized XY stage (Nikon), a Z-piezo stage insert (PiezoConcept), a large incubator box for temperature control (OkoLab), and a small imaging chamber for CO_2_ control (OkoLab with custom adapters machined by the MPI-NAT’s machine shop). For all imaging experiments, the 100x CFI Plan Apo NA 1.45 (Nikon) objective was used. For the STED imaging, the image size was set to 25 µm x 25 µm with 50 nm pixel size (501 px x 501 px), a pixel dwell time of 10 µs with 2-line accumulation, and pinhole size 64 µm. 5 z-slices were acquired with 0.3 µm spacing. The 640 nm laser was used for excitation (at 12% power) and the 775 nm laser was used for STED depletion (at 7.45% power). The TTL communication was established using a NI-6008 (National Instruments) card.

### Software

NIS-Elements AR (v5.21, Nikon including the NIDAQ plug-in), the Stedycon software and a standard installation of ImageJ/FIJI run on a single Windows workstation was used to run the AutoSTED workflow. For subsequent image analysis scripts in Image/FIJI were used, the resulting data was then analyzed in Excel (Microsoft) and R (R Core Team), and figures were assembled in Affinity Designer (Affinity).

## Conflict of interest statement

The authors declare no competing financial interests.

## Data and availability

No new or unique reagents were generated in this study. Scripts are available at https://doi.org/10.17617/3.AW998E.

## Acknowledgements

We thank all members of the Lenart laboratory at MPI-NAT, in particular Jasmin Jakobi, Antonio Politi for protocols, reagents and support, and Monica Gobran for careful reading of the manuscript. M.I. is a member and has been funded by the IMPRS Molecular Biology Programme. The research group of P. L. is funded by the Max Planck Society.

## Author contributions

D.L., M.I. and P.L. designed and conceptualized the experiments. D.L. and M.I. conducted experiments and analyzed data. P.L. analyzed data and wrote the manuscript. D.L. and M.I. revised and improved the manuscript.

## Figure legends

**Supplemental Figure S1**. A photograph of the setup.

## Notes

https://doi.org/10.17617/3.AW998E

### Competing Interest Statement

The authors have declared no competing interest.

