## Supplementary figures and images for "AutoSTED: An automated workflow for STED super-resolution imaging of cell nuclei"

### Supplemental Figure S1

**Figure S1.**

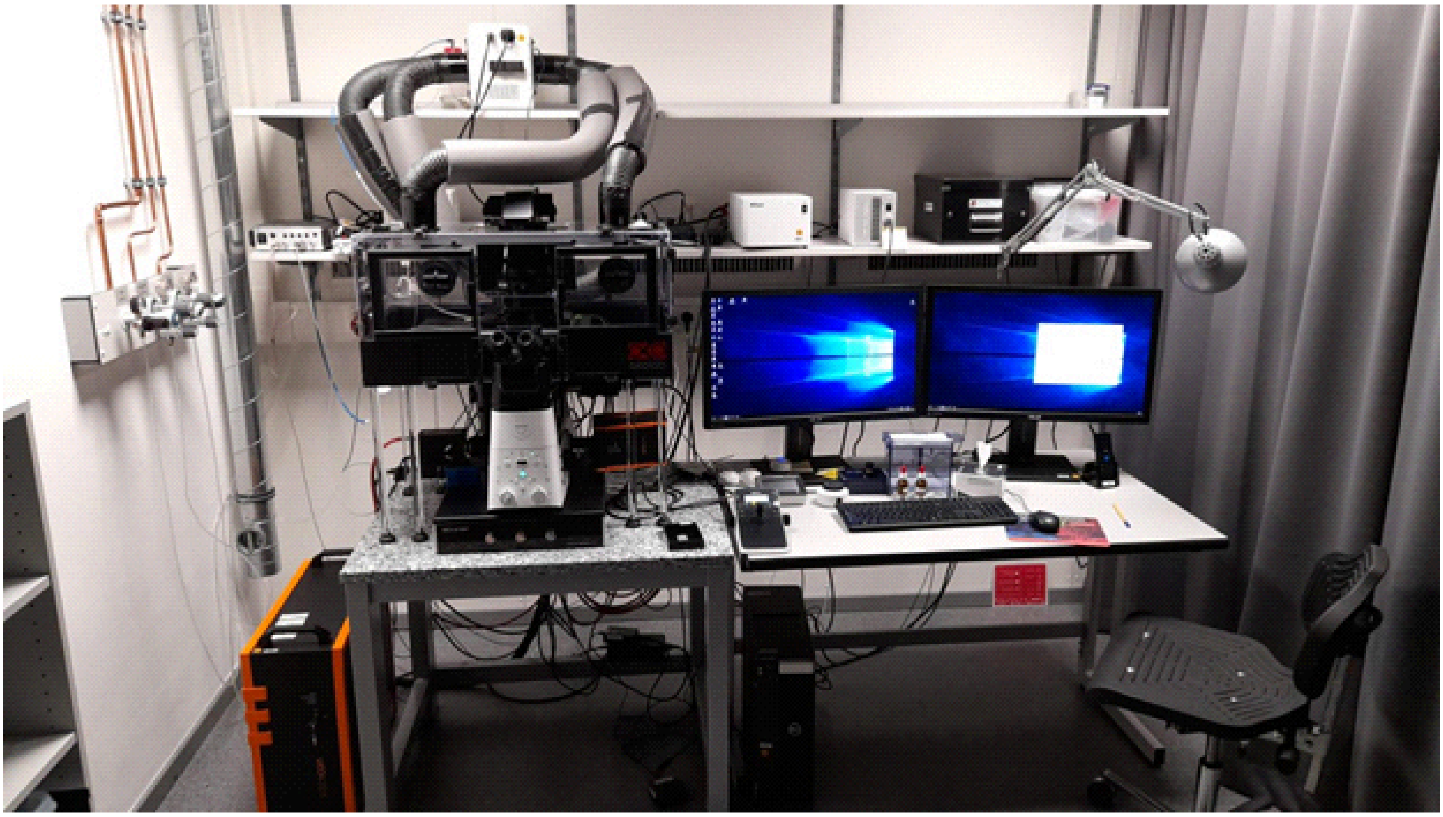
